# Loss of *Ythdf3* causes Danon disease-like features

**DOI:** 10.1101/2025.06.10.658803

**Authors:** Cong Yu, Chenzhong Xu, Jie Zhang, Ming Wang, Minxian Qian, Baohua Liu

## Abstract

YTH domain-containing family (YTHDF1/2/3) recognizes m6A modified mRNAs and regulates their stability, translation and function. We found loss of *Ythdf3* in mice caused cardiac hypotrophy, myopathy, and intellectual abnormalities, resembling the clinical features of a rare inherited X-linked disorder Danon disease (DD). Mechanistically, this was attributable to the compromised mRNA decay of sex-determining region Y (SRY)-box 9 (*Sox9*), mediated by YTHDF3 m6A reader function. Targeted therapy with AAV-shRNA against *Sox9* ameliorated fibrosis, increased neuron number, and significantly improved heart and brain function in *Ythdf3*^−/−^ mice. Our data reveal that loss of murine *Ythdf3* recapitulates systemic DD-like features, attributable to impaired *Sox9* decay, and highlight a novel therapeutic target for DD.

## Introduction

Danon disease (DD) is a rare inherited X-linked disorder, characterized by hypertrophic cardiomyopathy, myopathy, intellectual disability ^1,2,3^. Mutations of *LAMP2*, which encodes lysosomal associated membrane protein 2, were identified as the predominant causes of DD ^4^. During the last two decades, though DD pathobiology has been studied extensively, and heart transplantation is still the only therapeutic choice for DD at the end stage ^5^. LAMP2B is one of the three alternate splicing variants of *LAMP2* ^6^. Recent study showed that the treatment with AAV9-*LAMP2B* viral particle reversed autophagic flux defects and restored heart function in *Lamp2* deficient mice; however, brain defect was barely rescued ^7^. Therefore, understanding the underlying molecular mechanism of DD is still necessary.

Posttranscriptional modifications ensure a rapid response to environmental changes to maintain tissue/organ homeostasis ^8^. N6-methyladenosine (m^6^A) modification is one of the most prevalent posttranscriptional modifications in eukaryotic mRNAs, mainly catalyzed by methyltransferase-like 3/14 (METTL3/14), and Wilms tumor 1-associated protein (WTAP) (writers), and removed by demethylase fat mass and obesity-associated protein (FTO) and alkB homolog 5 (ALKBH5) (erasers) ^9^. The m6A modified mRNAs are recognized by a broad array of RNA-binding reader proteins, such as YTH domain-containing proteins (YTHDFs), thus regulating the stability, translation and function ^9^. Extensive studies have established the significance of m6A modifications in regulating diverse physiological functions. Loss of *Mettl3* or *Mettl14* causes embryonic lethality; their tissue specific absence causes cardiac dysfunction, osteoporosis, liver fibrosis, immunodeficiency, delayed neurogenesis, and abnormal skeletal muscle development ^10-15^. Similarly, *Ythdf2*^−/−^ mice are embryonic lethal, while tissue-specific depletion of *Ythdf2* causes cardiac malfunction, alters muscle size, and perturbs neurodevelopmental trajectories ^16-18^. We have recently found that loss of *Ythdf1* accelerates murine aging; mechanistically, YTHDF1 anchors tuberous sclerosis complex 2 (TSC2) to lysosome surface via LAMP2B, thus inhibiting mTORC1 and delaying aging ^19^. In this study, we generated *Ythdf3* knockout (KO) mice to investigate the systemic impacts. Interestingly, *Ythdf3* KO mice exhibited DD-like phenotypes and multiple tissue fibrosis. At the molecular level, YTHDF3 promoted *Sox9* mRNA decay and targeted *Sox9* knockdown ameliorated fibrosis, and DD-like features in *Ythdf3*^−/−^ mice.

## Results

### Loss of *Ythdf3* causes Danon disease-like features in mice

To investigate the physiological role of Ythdf3, we generated a murine *Ythdf3* KO allele (**Fig. S1a**). Successful deletion of *Ythdf3* was validated by RT-PCR and western blotting (**Fig. S1b, c**). Next, we executed systematic analysis of *Ythdf3*^−/−^ mice at 3 months (m) old. An enlargement of the cardiac chamber and a significant increase in cardiomyocyte size was noticed in male *Ythdf3*^−/−^ mice compared to *Ythdf3*^+/+^ control mice (**Fig. 1a, b**). Echocardiography revealed cardiac dysfunction in male *Ythdf3*^−/−^ mice, as evidenced by reduced left ventricle ejection fraction (EF) and fractional shortening (FS) and increased LV trace volume (**Fig. 1c**). By contrast, such changes were not observed in age-matched female *Ythdf3*^−/−^ mice (**Fig. S2a-c**). Further, the brain size of male *Ythdf3*^−/−^ mice was obviously reduced compared with *Ythdf3*^+/+^ mice (**Fig. 1d**). Nissi staining and periodic acid-Schiff (PAS) staining indicated neuronal reduction and glycoprotein aggregation in the male *Ythdf3*^−/−^ brain compared with that of *Ythdf3*^+/+^ mice (**Fig. 1e**). The Morris water maze assay showed *Ythdf3* deficiency compromised the short-term memory (24 h) of male mice, as evidenced by reduced activity in the target area and decreased residence time in the fourth quadrant and the platform, but not long-term memory (72 h) (**Fig. 1f, g**). Moreover, a decrease in myofiber size of skeletal muscle was noticed in male *Ythdf3*^−/−^ mice (**Fig. 1h**). Accumulation of acetylcholine esterase (AChE) was also observed in the skeletal muscle of male *Ythdf3*^−/−^ mice (**Fig. 1i**). Consistent with this, male *Ythdf3*^−/−^ mice had weaker running endurance (**Fig. 1j**). Of note, these histological changes and functional decline were not observed in female *Ythdf3*^−/−^ mice (**Fig. S2d-l**). Thus, *Ythdf3* deficiency caused cardiac hypertrophy, mental retardation, and skeletal muscle weakness specifically in male mice. Notably, the overt phenotypes of *Ythdf3*^−/−^ mice resembles human Danon disease (DD), a rare inherited X-linked disorder mainly affecting boys, and is characterized by hypertrophic cardiomyopathy, myopathy, intellectual disability^1,2,3^.

**Figure 1.**
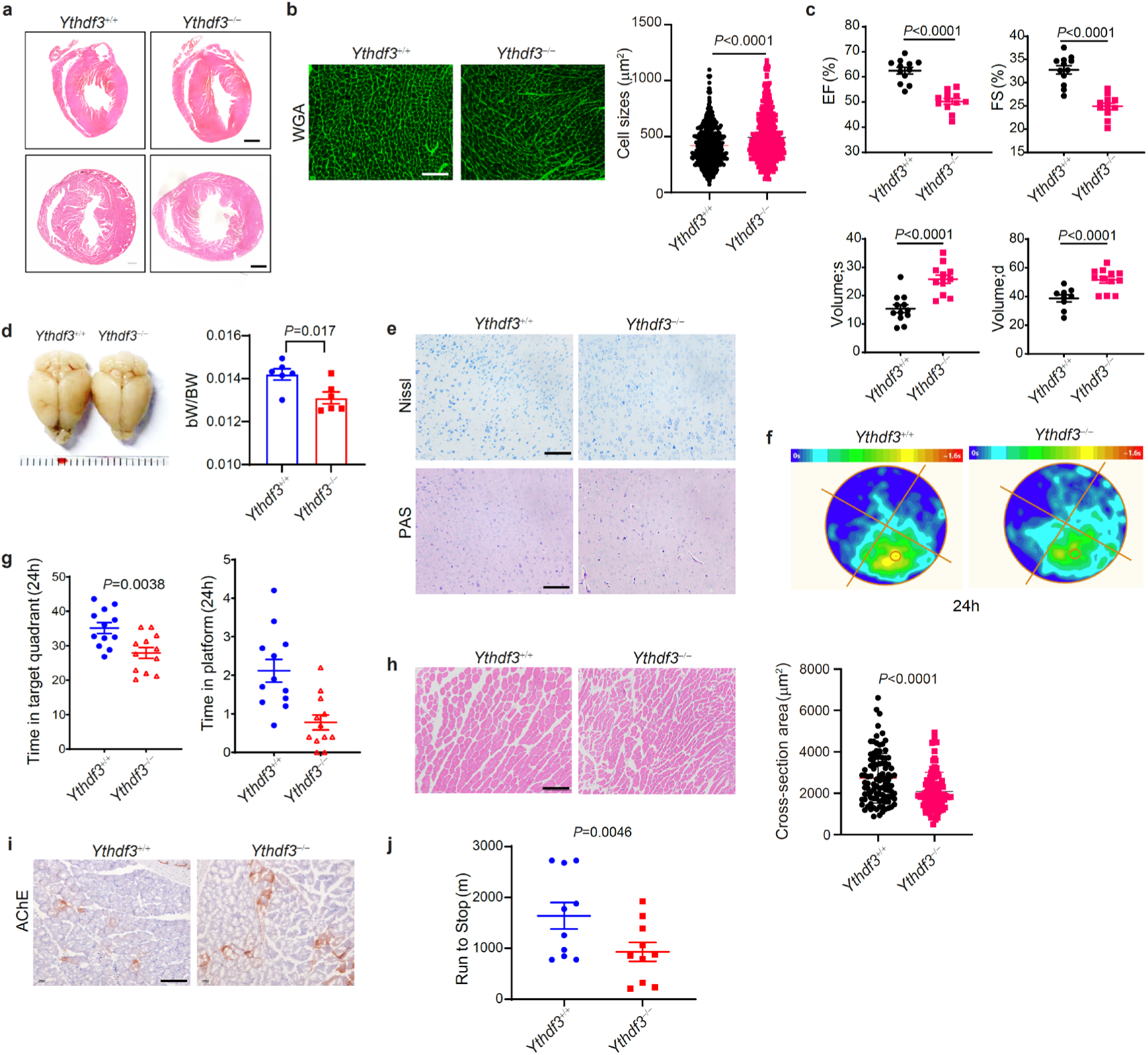
Loss of *Ythdf3* causes DD-like phenotype in mice a, Representative images of H&E-stained heart sections of male *Ythdf3^+/+^* and *Ythdf3*^−/−^ mice at 3 months (m) of age. Scale bar, 1 mm. **b,** Representative images of wheat germ agglutinin (WGA)-stained left ventricular muscle of male *Ythdf3^+/+^* and *Ythdf3*^−/−^ mice aged 3 m. Scale bar, 200 μm. Right: quantification of cell size, as determined by WGA staining. 500 cells from 5 mice per group were counted. **c,** Echocardiographic analysis showing left ventricle ejection fraction (EF), fractional shortening (FS) and left ventricular (LV) volume (s: systolic; d: diastolic) of male *Ythdf3^+/+^*and *Ythdf3*^−/−^ mice aged 3 m (n = 12). **d,** Left, representative images illustrating gross morphology of brains. Right, quantification of brain weight (bW)/body weight (BW) of male mice aged 3 m (n = 6). **e,** Representative images of Nissl-stained brain sections from male *Ythdf3^+/+^* and *Ythdf3*^−/−^ mice aged 3 m (top). Scale bar, 200 μm. Representative mages of periodic acid–Schiff (PAS)-stained brain sections from male *Ythdf3^+/+^* and *Ythdf3*^−/−^ mice aged 3 m (bottom). Scale bar, 200 μm. **f,** Heatmap of activity of mice in the Morris water maze 24 h post-training (n = 12 per genotype). **g,** Time mice spent in the quadrant and platform in the Morris water maze 24 h post-training (n = 12 per genotype). **h,** Representative images of H&E-stained skeletal muscle sections from male *Ythdf3^+/+^* and *Ythdf3*^−/−^ mice aged 3 m (top). Scale bar, 200 μm. Quantification of cross-section area of H&E staining (100 cells from 6 mice per group were counted) (bottom). **i,** Representative images of acetylcholinesterase (AChE) staining in skeletal muscle sections in male *Ythdf3^+/+^* and *Ythdf3*^−/−^ mice aged 3 m. Scale bar, 200 μm. **j,** Running endurance of male *Ythdf3^+/+^*and *Ythdf3*^−/−^ mice aged 3 m, as tested via treadmill (n = 10). Data represent the means ± SD. *P*-values were calculated by two-tailed unpaired Student’s *t*-test.

**Figure S1.**
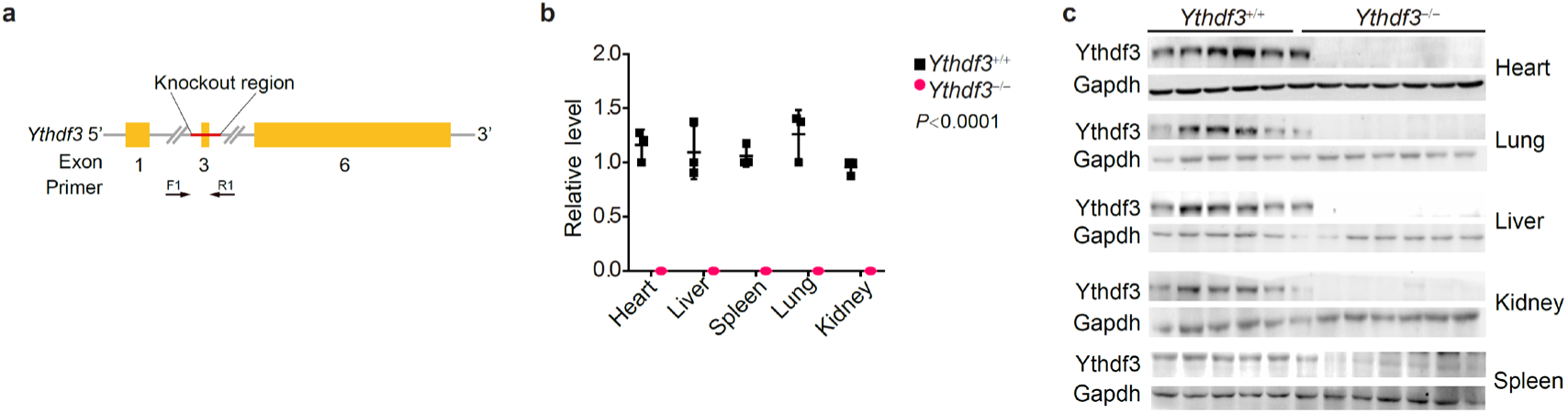
Generation of murine *Ythdf3* null allele. **a**, Strategy for generating *Ythdf3* knockout (KO) allele. F1 and R1: genotyping primers **b**, Quantitative PCR analysis of *Ythdf3* mRNA levels in tissues from male *Ythdf3^+/+^* and *Ythdf3*^−/−^ mice (n = 3). **c**, Representative immunoblots showing expression of YTHDF3 in different tissues from male *Ythdf3^+/+^*and *Ythdf3*^−/−^ mice.

**Figure S2.**
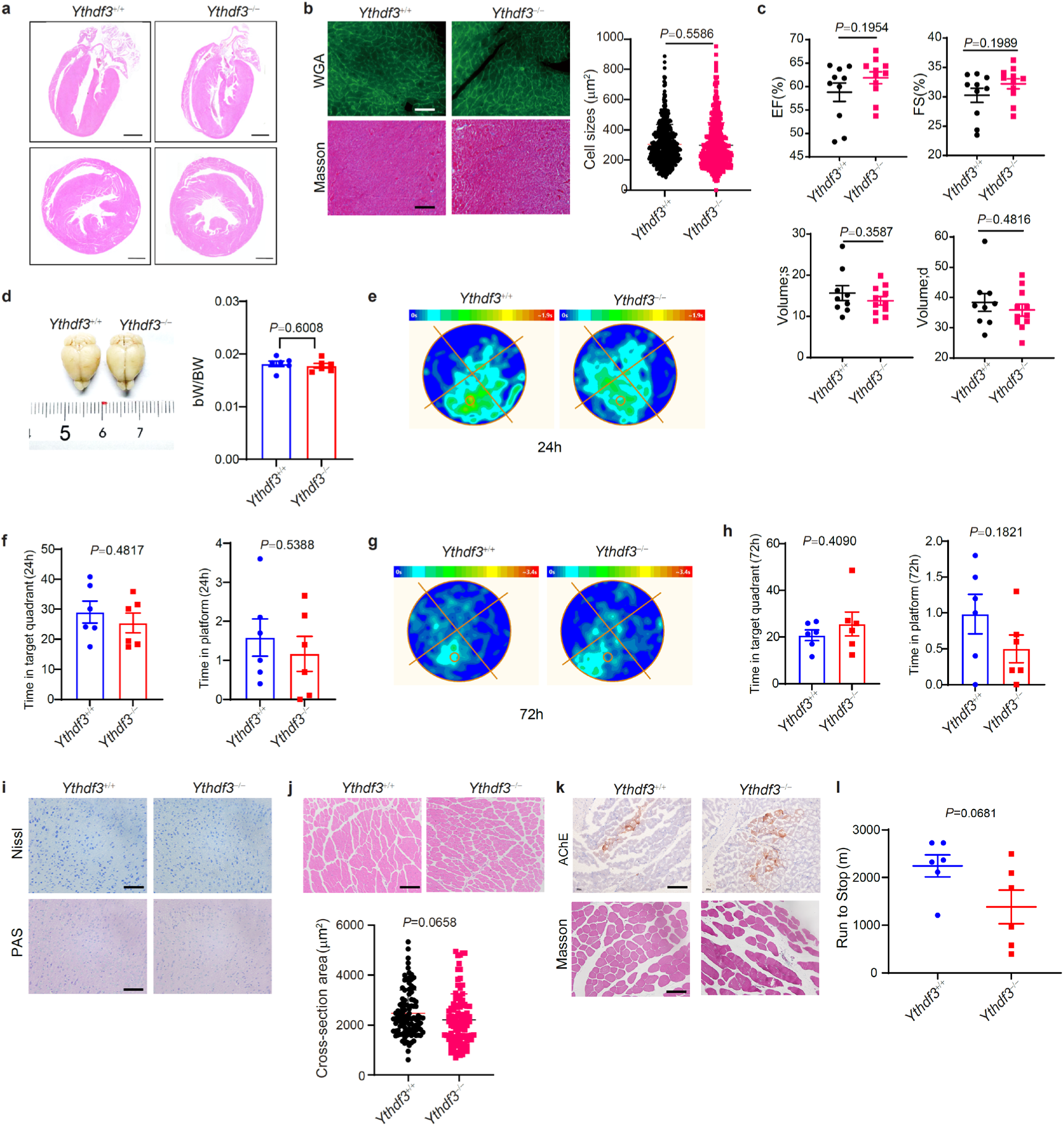
Characterization of female *Ythdf3*^−/−^ mice. **a**, Representative images of H&E-stained heart sections of 3-m-old female *Ythdf3^+/+^*and *Ythdf3*^−/−^ mice. Scale bar, 1 mm. **b**, Top: representative images of WGA-stained left ventricular muscle sections of 3-m-old female *Ythdf3^+/+^* and *Ythdf3*^−/−^ mice. Bottom: representative images of Masson’s trichrome-stained heart sections from 3-m-old female *Ythdf3^+/+^* and *Ythdf3*^−/−^ mice. Scale bar, 200 μm. Right: quantification of cardiac cell size in WGA-stained sections; 500 cells from 5 mice per group were counted. **c**, Echocardiography data of 3-m-old female *Ythdf3^+/+^* and *Ythdf3*^−/−^ mice (n = 10-11). **d**, Left, representative images of murine brains. Right, brain weight (bW) and body weight (BW) of female mice aged 3 m (n = 6). **e**, Heatmap of activity of female mice in the Morris water maze 24 h after training (n = 6). **f**, Time female mice spent in the quadrant and platform in the Morris water maze 24 h after training (n = 6). **g**, Heatmap of activity of female mice in the Morris water maze 72 h after training (n = 6). **h,** Time female mice spent in the quadrant and platform in the Morris water maze 72 h after training (n = 6). **i,** Top: representative images of Nissl-stained brain sections of 3-m-old female *Ythdf3^+/+^*and *Ythdf3*^−/−^ mice. Bottom: representative mages of periodic acid–Schiff (PAS)-stained brain sections from 3-m-old female *Ythdf3^+/+^*and *Ythdf3*^−/−^ mice. Scale bar, 200 μm. **j**, Top: representative images of H&E-stained skeletal muscle sections from 3-m-old female *Ythdf3^+/+^* and *Ythdf3*^−/−^ mice. Scale bar, 200 μm. Bottom: quantification of cross-section area of H&E staining (100 cells from 6 mice per group were counted). **k**, Top: representative images of acetylcholinesterase (AChE) staining in skeletal muscle sections from 3-m-old female *Ythdf3^+/+^* and *Ythdf3*^−/−^ mice. Bottom: representative images of Masson’s trichrome-stained skeletal muscle sections from 3-m-old female *Ythdf3^+/+^* and *Ythdf3*^−/−^ mice. Scale bar, 200 μm. **l**, Running endurance of 3-m-old female *Ythdf3^+/+^*and *Ythdf3*^−/−^ mice, as tested via treadmill (n = 6). Data represent the means ± SD. *P*-values were calculated by two-tailed unpaired Student’s *t*-test.

### YTHDF3 is a key suppressor of tissue fibrosis

A systemic Mason’s trichrome staining showed increased interstitial fibrosis in multiple tissues (heart, liver, muscle, and kidney) of 3-m-old male *Ythdf3*^−/−^ mice (**Fig. 2a**). α-smooth muscle actin (α-SMA) levels were elevated in examined tissues (**Fig. 2b**). Collagen I and Ⅲ levels were elevated in heart tissues of 3-m-old *Ythdf3*^−/−^ mice (**Fig. 2c**), affirming cardiac fibrosis. We focused on hearts for mechanistic investigation. To that end, we cultured *Ythdf3*^+/+^ and *Ythdf3*^−/−^ cardiac fibroblasts and reconstituted them with *Ythdf3* or *Ythdf3* m6A-reader-deficient mutant (*Ythdf3*-mut). Depletion of *Ythdf3* significantly upregulated gene expression of *FN1*, *COL1A1*, and *COL3A1*, while overexpression of *Ythdf3* downregulated them in both *Ythdf3*^−/−^ and *Ythdf3*^+/+^ cells. Of note, this effect was completely abolished in case that *Ythdf3*-mut was overexpressed (**Fig. 2d**).

**Figure 2.**
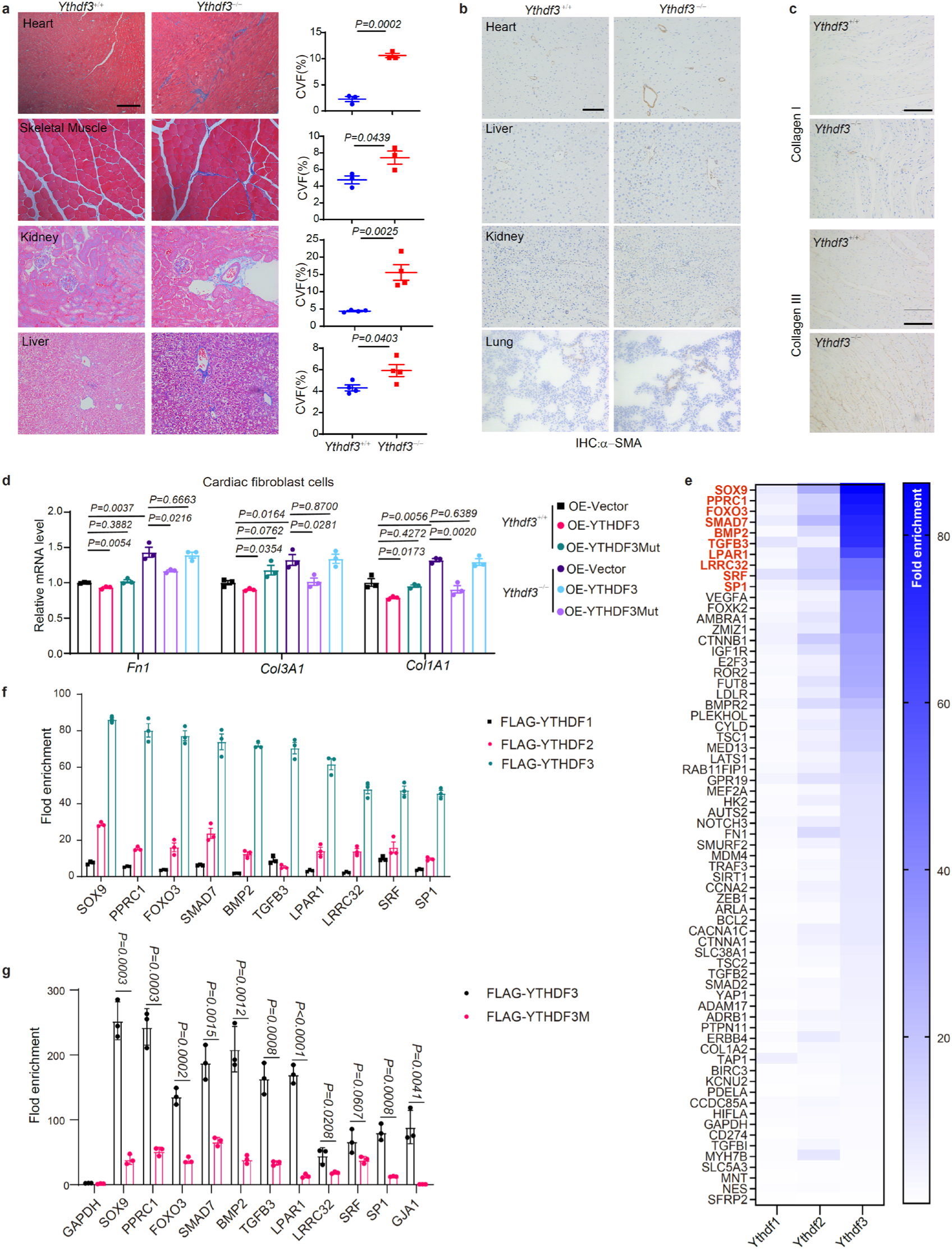
Loss of *Ythdf3* causes tissue fibrosis in male mice. **a**, Representative images of Masson’s trichrome-staining of heart, liver, muscle, kidney sections of 3-m-old male *Ythdf3^+/+^* and *Ythdf3*^−/−^ mice (left). Quantification of collagen-volume fraction (CVF%) in left (right). **b**, Representative images of immunostaining of α-SMA in heart, liver, kidney, and lung sections of 3-m-old male *Ythdf3^+/+^* and *Ythdf3*^−/−^ mice. **c**, Representative images of immunohistochemistry of Collagen I and Ⅲ in heart sections of 3-m-old male *Ythdf3^+/+^* and *Ythdf3*^−/−^ mice. **d**, Quantitative PCR analysis of collagen-related fibrosis genes in cardiac fibroblasts (P4) treated with vector, FLAG-YTHDF3 and -YTHDF3Mut (W438A and W492A mutations) lentivirus particles (n = 3 biological replicates). **e-f**, mRNA levels of the 67 genes that bound to YTHDFs (**Table S1** and **Fig. S4a**), determined by qRT-PCR in the anti-FLAG RIP immunoprecipitates in HEK293T cells. (**e**) Heatmap depicting relative levels of mRNAs bound to YTHDFs. (**f**) RIP-PCR showing top 10 mRNAs with the strongest binding capacity to YTHDF3 (n = 3 biological replicates). **g**, RIP-PCR analysis of mRNAs from **f** in HEK293T cells overexpressing FLAG-YTHDF3 and -YTHDF3M (W438A and W492A). *GJA1* was included as positive control ^23^, and *GAPDH* was included as negative control. n = 3 biological replicates. Data represent the means ± SD. *P*-values were calculated by two-tailed unpaired Student’s *t*-test. Scale bar, 200 μm.

**Figure S3.**
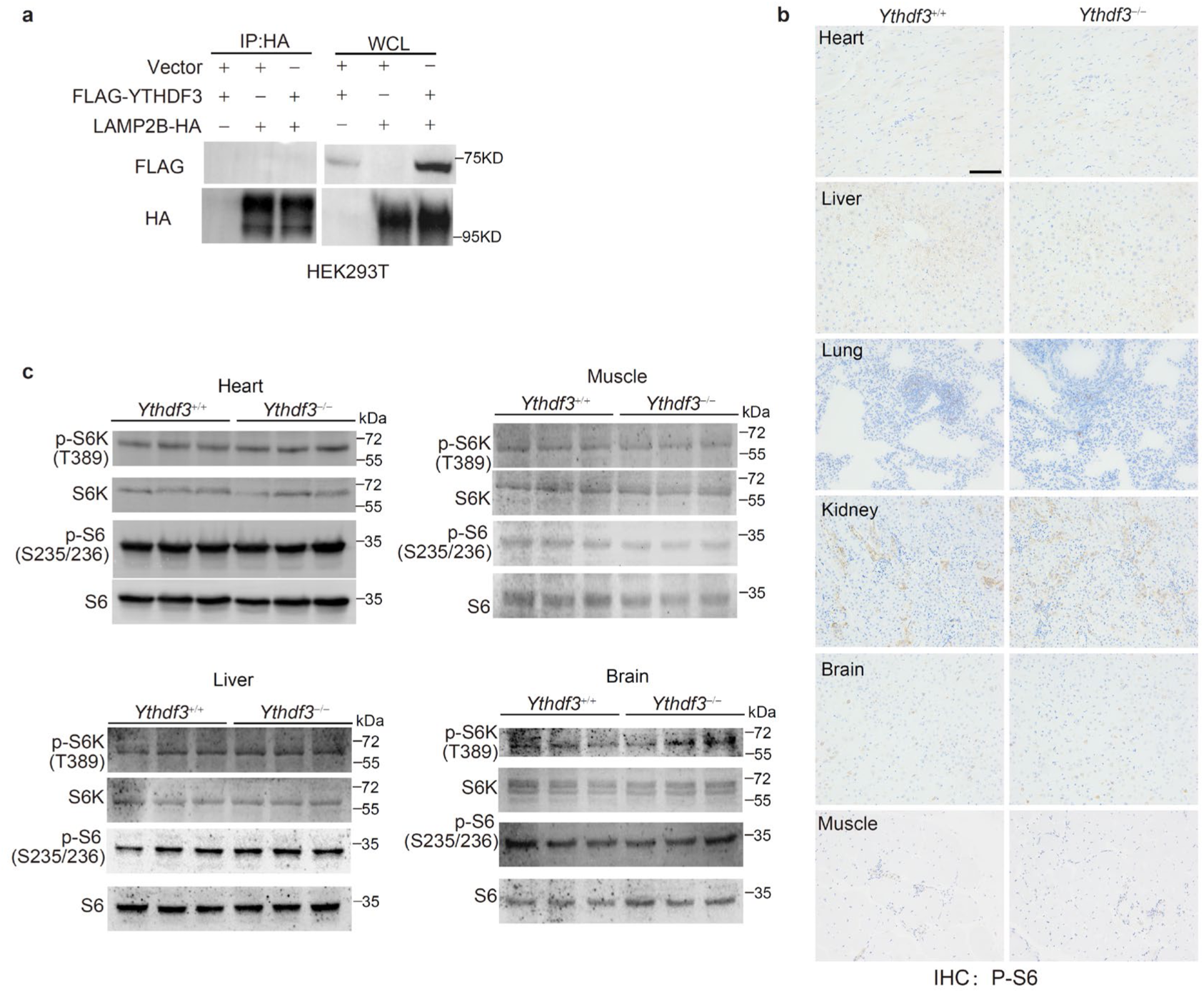
Loss of *Ythdf3* doesn’t activate the mTORC1 pathway a, Representative immunoblots showing proteins in anti-HA immunoprecipitates and whole-cell lysate (WCL) of HEK293T cells transfected with FLAG-YTHDF3 and/or LAMP2B-HA. **b,** Representative images of immunohistochemical staining (IHC) of p-S6 in tissues from 3-m-old male *Ythdf3^+/+^*and *Ythdf3*^−/−^ mice. Scale bar, 200 μm. **c,** Representative immunoblots showing protein expression in tissues from male *Ythdf3^+/+^* and *Ythdf3*^−/−^ mice aged 25 m.

**Figure S4.**
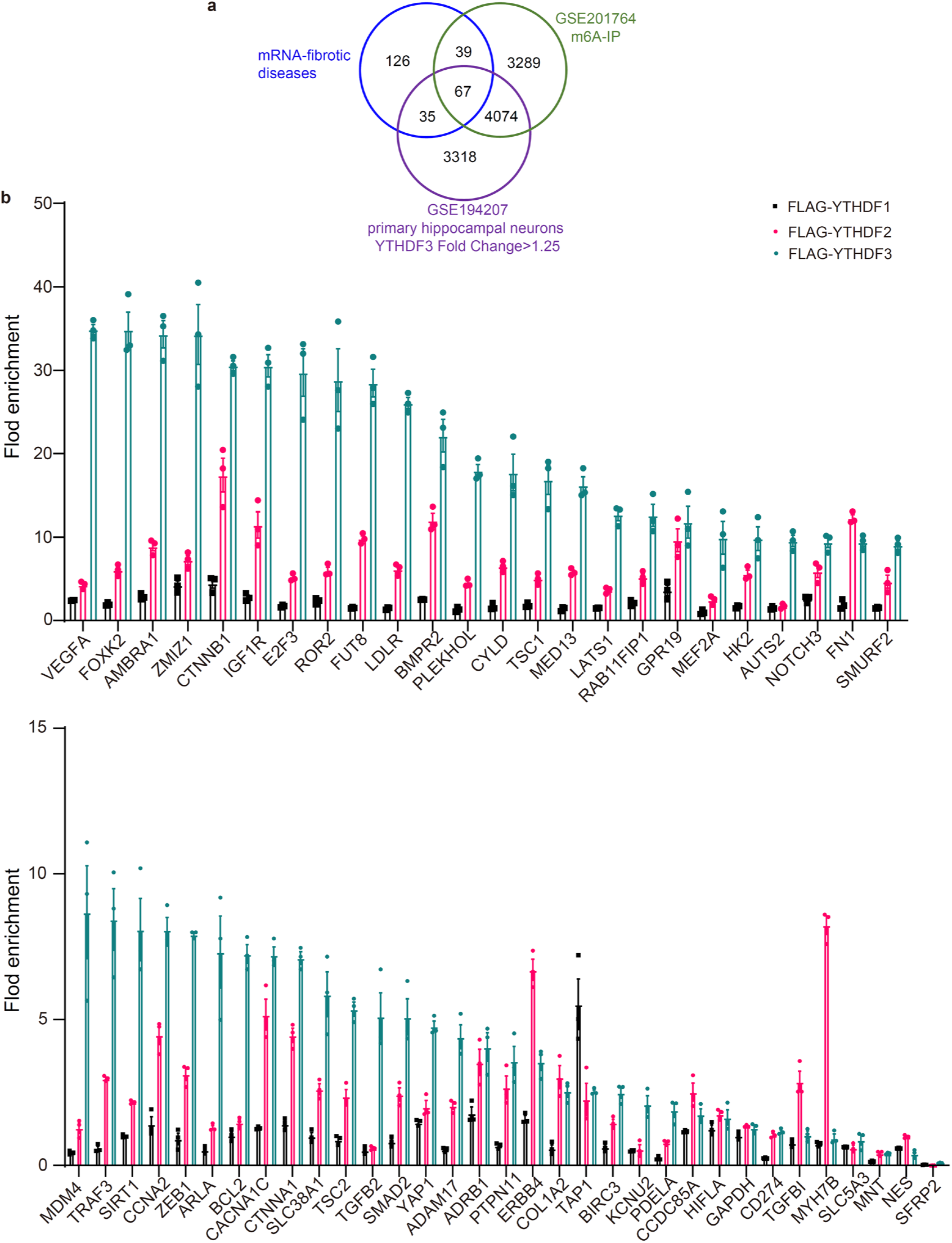
Fibrotic genes binding with YTHDFs. **a**, Venn diagram of the overlap among transcriptomes associated with fibrotic diseases (blue circle) ^22^, m6A-modified RNAs in mouse heart tissues (green circle, GSE201764), and YTHDF3-bound transcripts in murine primary hippocampal neurons (purple circle, GSE194207). **b**, Anti-FLAG RIP followed by quantitative PCR analysis of mRNAs in HEK293T cells overexpressing FLAG-YTHDFs (n = 3 biological replicates).

We next examined whether LAMP2B recruits YTHDF3 to inhibit mTORC1 activity in a similar manner to YTHDF1 ^19^. Co-immunoprecipitation (co-IP) was performed in HEK293T cells overexpressing FLAG-YTHDF3 and/or LAMP2B-HA. Interestingly, FLAG-YTHDF3 was hardly detected in the anti-HA immunoprecipitates (**Fig. S3a**). Moreover, mTORC1 hyperactivation of was not observed in *Ythdf3*^−/−^ mice (**Fig. S3b, c**), suggesting *Ythdf3* deficiency-driven DD features is independent of mTORC1.

Next, we investigated YTHDF3-read m6A-transcriptomes. We took advantage of the public database GSE201764 (identifying m6A-modified transcripts in mouse hearts) ^20^, and GSE194207 (identifying YTHDF3-bound mRNAs in mouse hippocampal neurons) ^21^, and searched for YTHDF3-read m6A-modified mRNAs that are associated with fibrotic diseases ^22^. A total of 67 fibrotic genes were enriched (**Table S1** and **Fig. S4a**), and their associations with YTHDFs were validated by RNA IP (RIP) with anti-FLAG antibody in HEK293T cells overexpressing FLAG-YTHDFs (**Fig. 2e, f**, and **Fig. S4b**). Of note, the top 10 mRNAs showing the strongest binding ability to YTHDF3 are *SOX9*, *PPRC1*, *FOXO3*, *SMAD7*, *BMP2*, *TGFB3*, *LPAR1*, *LRRC32*, *SRF*, and *SP1* (**Fig. 2f**). Their bindings to m6A-reader-deficient FLAG-YTHDF3M were completely abolished (**Fig. 2g**). These data suggest YTHDF3 as an important regulator of cardiac fibrosis by an m6A-dependent mechanism.

### YTHDF3 mediates *Sox9* mRNA decay

*SOX9*, which encodes a SOX family transcription factor involved in sex determination ^24^, is the top differential binding transcript with YTHDF3, and its abnormal upregulation induces cardiac fibrosis ^25^. Indeed, both mRNA and protein levels of SOX9 and its transcription target genes ^25^ were significantly increased in male *Ythdf3*^−/−^ hearts, as determined by quantitative PCR and immunofluorescence (IF) microscopy (**Fig. 3a-c**). Upregulation of Sox9 downstream genes was noticed in *in vitro* cultured *Ythdf3*^−/−^ cardiac fibroblasts, which was restored by overexpression of *Ythdf3* (**Fig. S5**). Considering the male specificity of DD features in *Ythdf3*^−/−^ mice, we focused on SOX9 for mechanistic investigation. *SOX9* mRNA undergoes m6A modification ^26^. *Sox9* mRNA degradation was significantly impaired in *Ythdf3*^-/-^ cardiac fibroblasts (**Fig. 3d**). We mutated the m6A sites on *Sox9* transcript (*SOX9m*). In HEK293T cells overexpressing *SOX9* or *SOX9m*, RIP results showed YTHDF3 interacted with the *SOX9* transcript in an m6A dependent manner (**Fig. 3e**). Further, an ectopic truncation of *Sox9* cDNA (a 2040-2500 bp m6A-rich sequence, t*Sox9*) or m6A mutated (t*Sox9m*) was overexpressed in murine cardiac fibroblast cells. A significant decrease in the degradation rate of t*Sox9m* mRNA was observed in both *Ythdf3*^+/+^ and *Ythdf3*^-/-^ cells (**Fig. 3f**). We subsequently generated an m6A-deficient Sox9 mutant (Sox9m) by introducing A-to-G mutations at all four m6A sites (positions 2184, 2163, 2104, and 2096 ^27^), and the subcellular distribution of *Sox9* and *Sox9m* transcripts was examined using the MS2 and MS2-coat protein (MCP) imaging system ^28^. While *Sox9* transcripts accumulated with YTHDF3, *Sox9m* transcripts were barely co-localized with YTHDF3 (**Fig. 3g**). This suggests that m6A modification is required for the association of YTHDF3 with *Sox9* mRNA. In line with this, knocking down *Mettl3* in mouse cardiac fibroblasts reduced YTHDF3’s binding to *Sox9* transcripts (**Fig. 3h, i**). Notably, protein degradation rate of SOX9 was not affected by *Ythdf3* deficiency (**Fig. 3j**). Thus, m6A reader function of YTHDF3 mediates *Sox9* mRNA decay.

**Figure 3.**
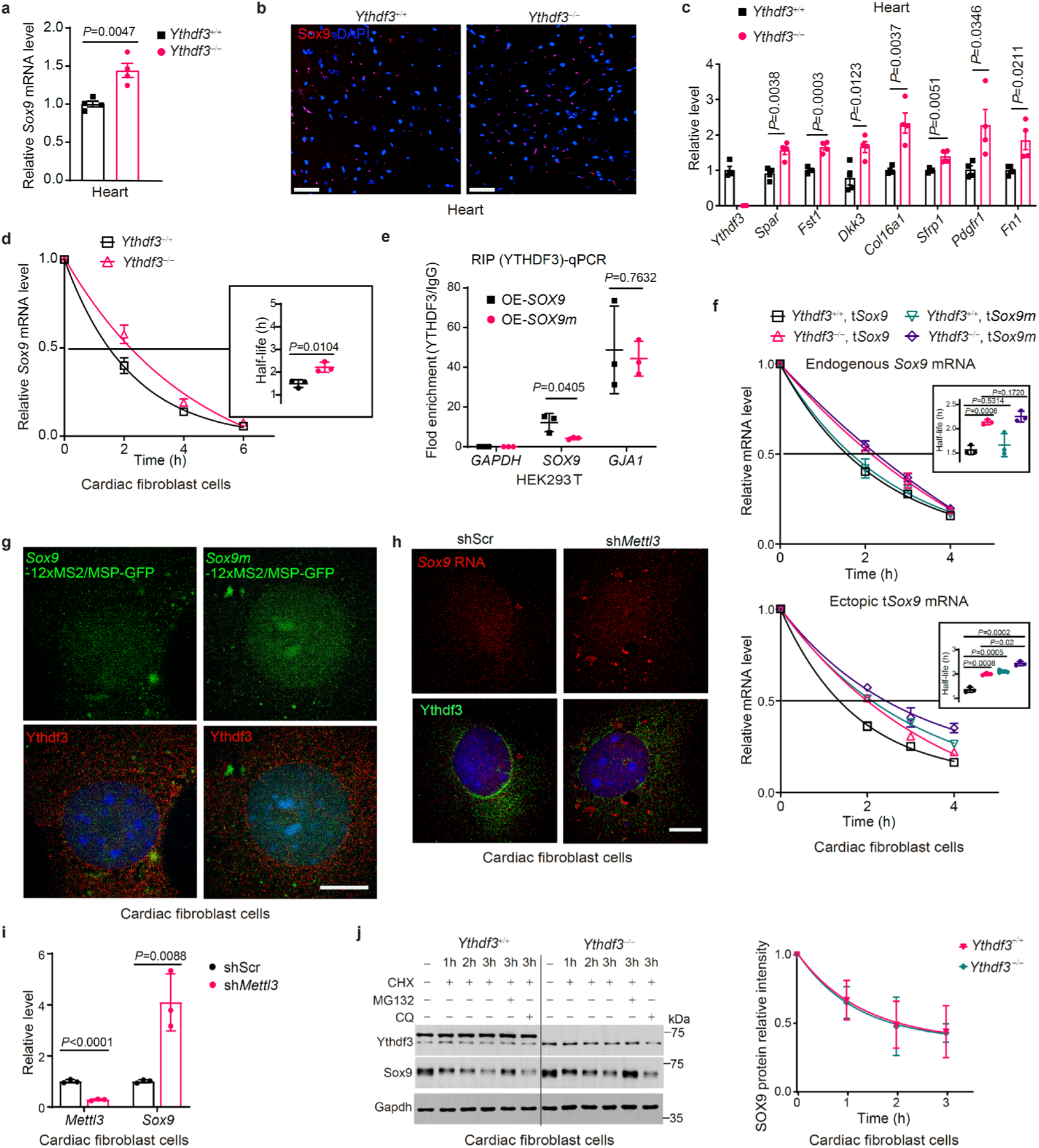
YTHDF3 regulates mRNA decay of *Sox9* in an m6A-dependent manner. **a**, Quantitative PCR analysis of *Sox9* mRNA levels in heart tissues of male *Ythdf3^+/+^* and *Ythdf3*^−/−^ mice aged 3 m. **b**, Representative fluorescence images showing Sox9 protein in heart tissues of 3-m-old male *Ythdf3^+/+^* and *Ythdf3*^−/−^ mice (n = 4). Scale bar, 30 μm. **c**, Quantitative PCR analysis of *Sox9* downstream gene levels in heart tissues of 3-m-old male *Ythdf3^+/+^* and *Ythdf3*^−/−^ mice (n = 4). **d**, *Sox9* mRNA degradation analysis in *Ythdf3^+/+^* and *Ythdf3*^−/−^ cardiac fibroblast. Insert: comparation of calculated half-life of *Sox9* mRNAs. **e**, YTHDF3-RIP followed by quantitative PCR analysis of *SOX9* and *SOX9 mutation* (m6A sites) mRNA levels in HEK293T cells overexpressing *SOX9* and its mutation (n = 3 biological replicates). **f**, Degradation analysis of endogenous *Sox9*, ectopic truncated *Sox9* (t*Sox9*) and its m6A mutation (t*Sox9m*) in male *Ythdf3^+/+^* and *Ythdf3*^−/−^ cardiac fibroblasts overexpressing the t*Sox9* or t*Sox9m* (n = 3 biological replicates). Left: detection of endogenous *Sox9*; right: detection of t*Sox9* and t*Sox9m*. Insert: half-life analysis of *Sox9* variants. **g**, Representative fluorescence images showing *Sox9*-, *Sox9* with m6A mutation (*Sox9m*)*-*12×MS2/MCP-EGFP and Ytthdf3 (red) in cardiac fibroblasts. Scale bar, 15 μm. **h**, Representative fluorescence images showing *Sox9* mRNA (Cy3) and Ythdf3 (green) in mouse cardiac fibroblasts transfected with sh*Scr* or sh*Mettl3*. Scale bar, 15 μm. **i**, Quantitative PCR analysis of *Sox9* mRNA levels in scramble (shScr) and sh*Mettl3* shRNA-transfected cardiac fibroblasts (n = 3 biological replicates). **j**, Representative immunoblots showing SOX9 protein level in cardiac fibroblasts from male *Ythdf3^+/+^* and *Ythdf3*^−/−^ mice after treatment with cycloheximide (CHX) (50 µg/mL); chloroquine (CQ, 50 μM); or MG132 (10 μM). Right: Quantification of the SOX9 protein level was shown. Data represent the means ± SD. *P*-values were calculated by two-tailed unpaired Student’s *t*-test.

**Figure S5.**
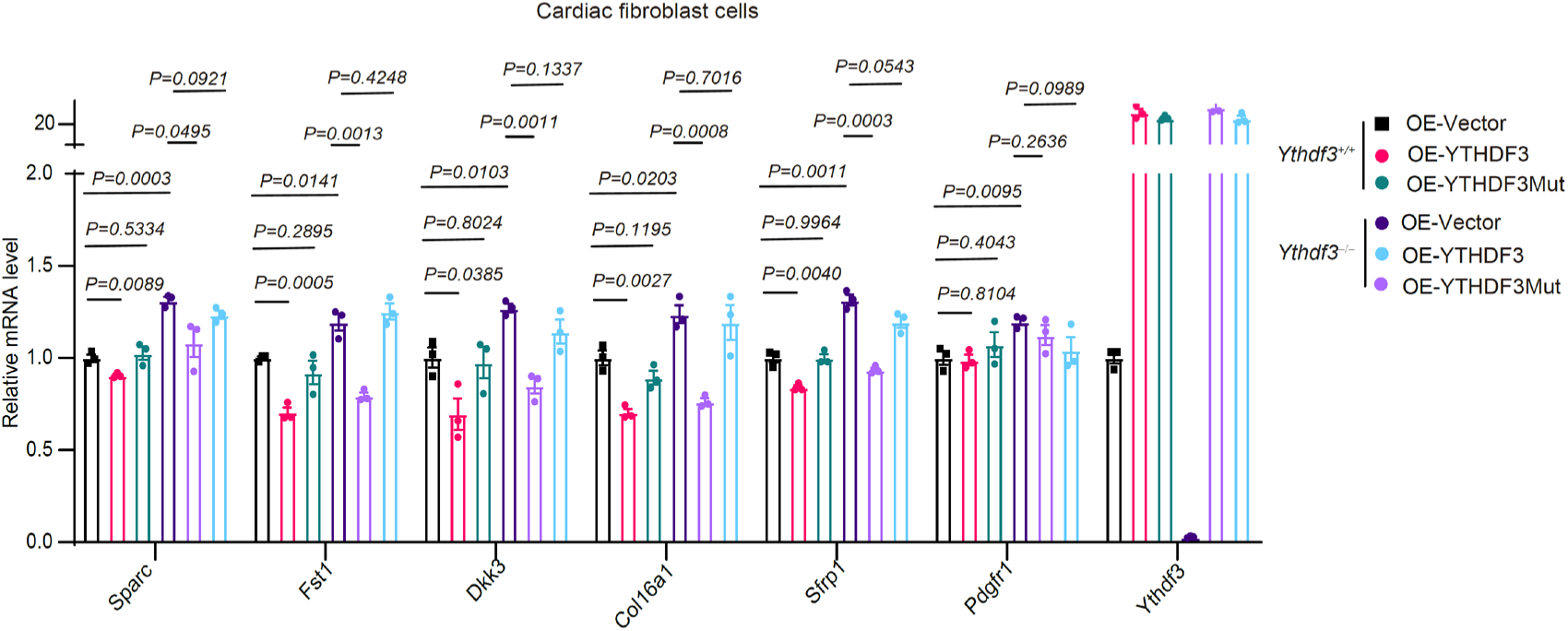
YTHDF3 regulates Sox9 target genes in an m6A-dependent manner. Quantitative PCR analysis of Sox9 downstream gene levels in *Ythdf3^+/+^* and *Ythdf3*^−/−^ cardiac fibroblast cells (P4) treated with vector, FLAG-YTHDF3 and -YTHDF3Mut (W438A and W492A) lentivirus particles (n = 3 biological replicates). Data represent the means ± SD. *P*-values were calculated by two-tailed unpaired Student’s *t*-test.

### Targeted inhibition of *Sox9* reverses DD features

Next, we employed an adeno-associated virus serotype 9 (AAV9) with a sh*Sox9* cassette (AAV-sh*Sox9*) to manipulate the level of SOX9 *in vivo* (**Fig. 4a**). Compared with mice given the scramble (AAV-Scr), *Ythdf3*^+/+^ and *Ythdf3*^-/-^ mice intravenously injected with AAV-sh*Sox9* showed downregulated mRNA and protein levels of SOX9 in hearts and brains (**Fig. 4b-d**). Markedly, the cardiac function of male *Ythdf3*^-/-^ mice was improved, reaching a level comparable to that of *Ythdf3*^+/+^ mice after AAV-sh*Sox9* therapy, as evidenced by an increase in EF and FS and a reduction in LV volume (**Fig. 4e, S6a**). AAV-sh*Sox9* therapy led to a significant decrease in the transcription levels of genes associated with cardiac fibrosis and alleviated cardiac fibrosis in male *Ythdf3*^-/-^ mice (**Fig. 4f, g**). An increase in neuronal count and a decrease in glycoprotein aggregation were evidenced after AAV-sh*Sox9* therapy (**Fig. 4i**). Consistent with this, the AAV-sh*Sox9* therapy improved the brain function of male *Ythdf3*^-/-^ mice, as in the Morris water maze tests performed 24 hours post-training, both the activity in the target area and residence time in the fourth quadrant increased in male *Ythdf3*^-/-^ mice given AAV-sh*Sox9* therapy (**Fig. 4h**, **S6b**). These findings suggest that targeting *Sox9* is beneficial for heart and brain health and function.

**Figure 4.**
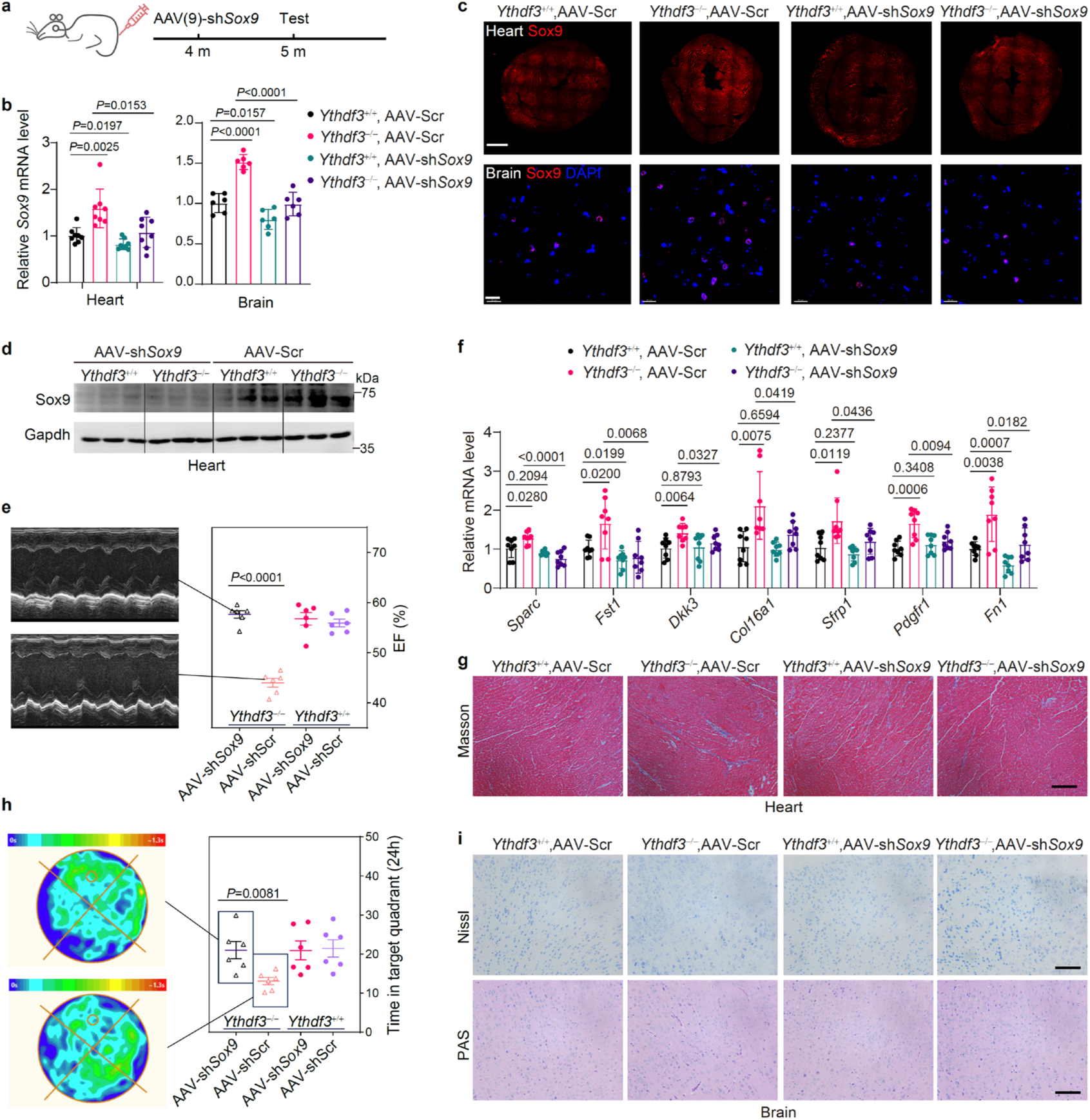
Targeted inhibition of *Sox9* ameliorates DD features in *Ythdf3*^-/-^ mice a, Design of AAV-sh*Sox9* therapeutic study. Injection was administered at 4 months and testing was conducted at 5 months. **b,** Quantitative PCR analysis of *Sox9* mRNA levels in heart and brain tissues after AAV-*shSox9* therapy. **c,** Representative fluorescence images showing SOX9 protein expression after AAV-*shSox9* therapy in heart (scale bar, 1 mm) and brain (scale bar, 20 μm) of *Ythdf3*^+/+^ and *Ythdf3*^-/-^ mice. **d,** Representative immunoblots showing SOX9 protein expression in heart tissues *Ythdf3*^+/+^ and *Ythdf3*^-/-^ mice after AAV-*shSox9* therapy. **e**, Echocardiographic analysis of *Ythdf3*^+/+^ and *Ythdf3*^-/-^ mice after AAV-*shSox9* therapy (n = 6). **f,** Quantitative PCR analysis of *Sox9* downstream gene levels in heart tissues of *Ythdf3*^+/+^ and *Ythdf3*^-/-^ mice after AAV-*shSox9* therapy (n = 8). **g,** Representative images of Masson’s trichrome staining of heart sections of *Ythdf3*^+/+^ and *Ythdf3*^-/-^ mice after AAV-*shSox9* therapy. Scale bar, 200 μm. **h,** Heatmap of activity (left) and time spent in target quadrant (right) of *Ythdf3*^+/+^ and *Ythdf3*^-/-^ mice given AAV-*shSox9* therapy in the Morris water maze 24 h after training (n = 6). **i,** Top: representative images of Nissl-stained brain sections from *Ythdf3*^+/+^ and *Ythdf3*^-/-^ mice after AAV-*shSox9* therapy. Bottom: representative images of periodic acid-Schiff (PAS)-stained brain sections from *Ythdf3*^+/+^ and *Ythdf3*^-/-^ mice after AAV-*shSox9* therapy. Scale bar, 200 μm. Data represent the means ± SD. *P*-values were calculated by two-tailed unpaired Student’s *t*-test.

**Figure S6.**
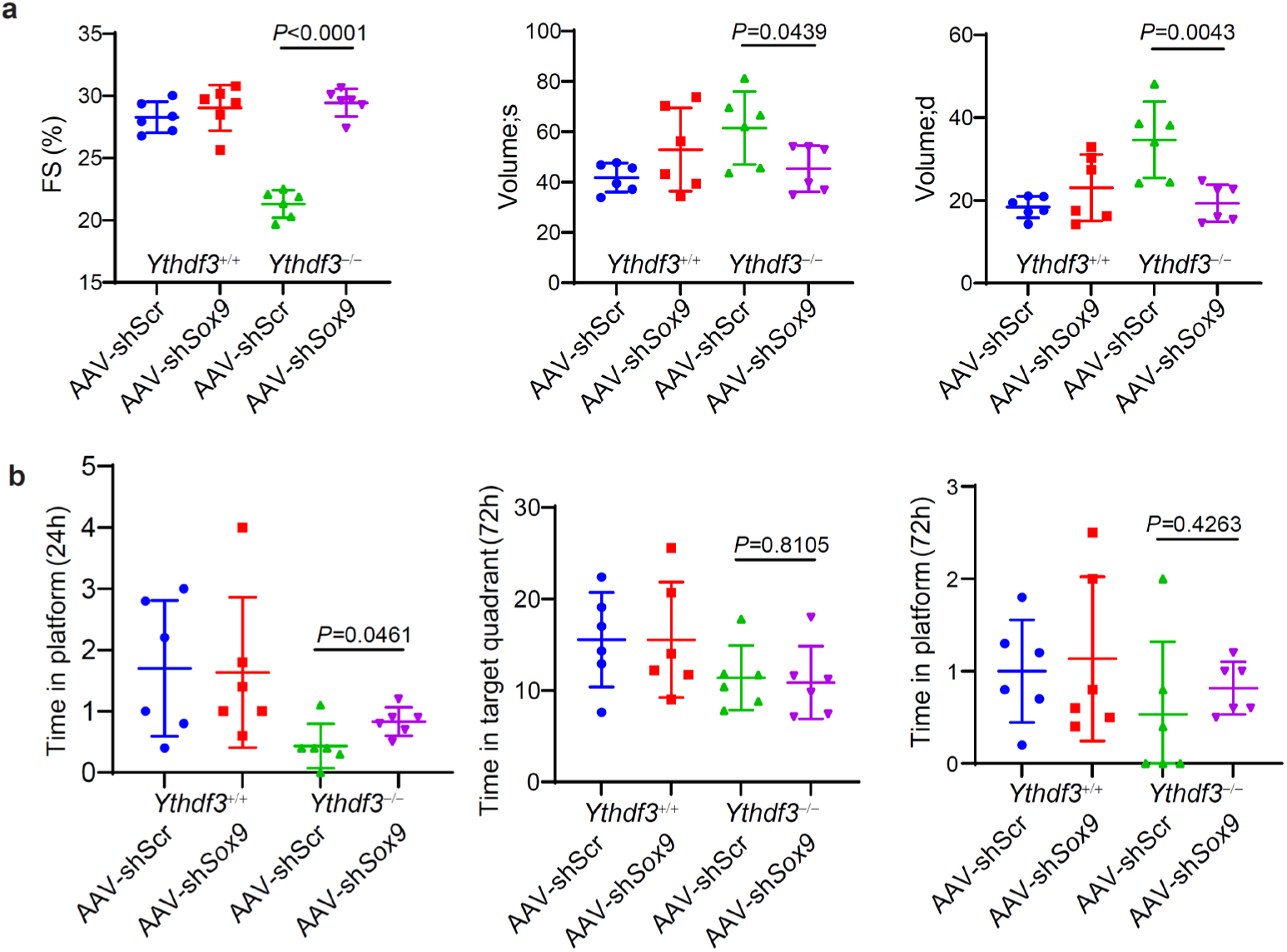
Targeting *Sox9* is beneficial for heart and brain function. **a**, Echocardiographic analysis showing fractional shortening (FS) and left ventricular (LV) volume (s: systolic, d: diastolic) in mice after AAV-*shSox9* therapy (n = 6). **b**, Time spent in platform (left) 24 h, and in target quadrant (middle) and platform (right) of mice given AAV-*shSox9* therapy in the Morris water maze 72 h after training (n = 6). Data represent the means ± SD. *P*-values were calculated by two-tailed unpaired Student’s *t*-test.

## Discussion

DD is predominantly caused by *LAMP2* mutations; however, the clinical treatment is restricted ^4,5^. In this study, we found that *Ythdf3* depletion induced DD-like features and multiple tissue fibrosis, attributable to compromised *Sox9* mRNA decay. AAV-mediated *Sox9* gene silencing ameliorates DD-related phenotypes in *Ythdf3*^-/-^ mice. Our findings provide a new DD mice model independent of LAMP2. It is necessary to study whether targeted *Sox9* gene therapy can ameliorate the DD features in *Lamp2* KO mice.

Fibrosis is a prevalent pathological feature, characterized by chronic inflammation, and excessive deposition of extracellular matrix (ECM) ^29^. m6A modification, its modifying enzymes and readers dynamically regulate tissue fibrosis. We found YTHDF3 mediates *Sox9* mRNA decay to suppress fibrosis. SOX9, a sex-determining gene, is essential for development, and declines after development ^30^. The appropriate temporospatial control of *SOX9* expression is critical for tissue homeostasis. Ectopic SOX9 expression beyond the developmental stages induces fibrosis ^31,32^, causing heart failure in adult mice^33^. In our study, the AAV-sh*Sox9* gene therapy completely restored the cardiac and neuronal defects in *Ythdf3* KO mice, suggesting potential of *SOX9* siRNA and inhibitor in clinical treatment of DD. Estrogen blocks the nuclear entry of SOX9, and thus its function^34,35^. This may explain why the clinical phenotypes of females with DD is often present later in life (usually 10 years later than men) and with isolated cardiac manifestations ^1^. Of note, we didn’t observe significant improvement in skeletal muscle after AAV-sh*Sox9* gene therapy, likely attributable to the limited efficacy of AAV delivery. Other delivery systems that target specific organs, such as exosomes or lipid nanoparticles (LNP), may be promising approaches in the future.

In summary, our evidence shows decline of YTHDF3 compromised *Sox9* mRNA decay, causing DD-like features, and targeting *Sox9* with shRNA inhibited the level of cardiac fibrosis in DD mice, thereby improving the heart and brain function. This not only aids in the understanding of the molecular pathogenesis of DD, but also holds great promise for the identification of novel therapeutic candidates for DD.

## Methods

### Reagents and antibodies

All reagents and antibodies are listed in **Table S2**. **Plasmids and constructs** LAMP2B-HA, FLAG-YTHDF3 were generated using PCR cloning in a pcDNA3.1 vector. The coding sequence of *Sox9* RNA was fused with the MS2 sequence. Subsequently, it was cloned into the pCDH-CMV-MCS vector by using a PCR cloning kit (C112-01; Vazyme, China), aiming to generate pCDH-Sox9-MS2. The MCP coding sequence and GFP were generated through PCR cloning into the pCDH-CMV-MCS vector to obtain MS2-GFP. All DNA site mutations are performed using a Mut Express MultiS Fast Mutagenesis kit (C215 -01; Vazyme, Nanjing, China). Short hairpin RNA (shRNA) sequences targeting specific genes (listed in **Table S3**) were cloned into the PLKO.1 vector using a DNA ligation kit (6023; TaKaRa, Kyoto, Japan). All constructs were verified by DNA sequencing (Ruibiotech, Guangzhou, China).

### Animals

*Ythdf3*-knockout (KO) alleles were created by CRISPR–Cas9-mediated genome editing in C57BL/6 mice by the transgenic animal services of Cyagen Biosciences (Suzhou, China). Briefly, exon 3 of *Ythdf3* was targeted with the following oligonucleotide sequences:

sgRNA1: 5′-AGTCACAAATAGTTACTTGAAGG-3′

sgRNA2: 5′-AAACATATACTGTGAAGCGTTGG3′

The Cas9 construct and gRNA were co-injected into fertilized eggs. The *Ythdf3* KO allele was detected using the following primers:

Forward: 5’-CTTCAGTGCATGCTAAATACAC-3’

Reverse: 5’-CTAAGATTTCAGACAATTTTCCAC-3’

*Ythdf3* wild-type allele was detected using the following primers:

Forward: 5’-GTTTTTATCTCCGTGTCTCTACTAG-3’

Reverse: 5’-CTAAGATTTCAGACAATTTTCCAC-3’

Mice were housed and handled in accordance with protocols (IACUC-202300033) approved by the Institutional Animal Care and Use Committee of Shenzhen University, China.

### Echocardiographic evaluation

Mice were anesthetized using 1.5-2% isoflurane gas inhalation, then subjected to transthoracic echocardiography (Vevo 2100 Imaging System, VisualSonics, USA). In brief, each animal was placed on a heated table in a supine position with the extremities fixed to the table with sellotape, and hair was depleted from the chest using a chemical hair remover (Veet, France). Heated ultrasound transmission gel (TM100, Jinya, Tianjin, China) was added to the surface of the thoracic cavity to optimize the visibility of the cardiac chamber. Heart rate (HR) was maintained at a consistent 475 ± 50 bpm in all experimental groups, and the core temperature was maintained at 37°C. Two-dimensional left ventricular B-mode images were acquired at the level of the papillary muscles in the parasternal long-axis view. Long-axis B-mode echocardiographic images of hearts during diastole and systole were obtained. Vevo 2100 software was used to calculate the percentage EF and FS, and other measurements, for each heart.

### Morris water maze

The Morris water maze assay was conducted as previously described^36^. In brief, mice were trained in a round water-filled tub (160 cm in diameter and 50 cm in height). Each mouse was given 4 trials (1 min) per day for 5 consecutive days. If mice did not reach the platform in the allotted time, they were manually guided to it, and kept there for 1 min. After a 5-day training period, time that mice spent in the original platform quadrant was recorded to assess the mice’s short-term (24 h) and long-term (72 h) memory consolidation. Data were recorded using an HVS water maze program (WaterMaze3, Actimetrics, Wilmette, IL, USA).

### Running endurance test

Running endurance was tested with treadmill apparatus (Sansbio, Nanjing, China), as described previously^37^. In brief, mice were trained for 5 min at a speed of 15 m/min on the treadmill for 2 consecutive days, then tested for 5 min at 15 m/min, 5 min at 20 m/min, 10 min at 25 m/min, and 30 m/min until failure.

### Gene therapy

Recombinant adeno-associated virus 9 (rAAV9)-overexpressing sh*Sox9* was obtained through a commercial service offered by WZ Biosciences (Jinan, China). In brief, the sh*Sox9* sequences (**Table S3**) were cloned into the pAV-U6-shRNA-CMV-GFP vector. The sh*Sox9* (1 × 10^12^ vg) virus and control virus (1 × 10^11^ vg) particles were injected into mice via the tail vein.

### Tissue section staining

Paraffin-embedded sections of PFA-fixed tissues were dewaxed, hydrated, and stained with H&E stain (G1076; Servicebio, Wuhan, China), Nissl stain (G1036; Servicebio), PAS stain (G1008; Servicebio), Masson’s trichrome stain (G1340; Solarbio, Beijing, China), and immunohistochemical stain (GK600710; Genetech, Shanghai, China) with specific antibodies, following the manufacturer’s instructions. Sections were mounted with neutral balsam mounting medium (E675007; BBI Life Science, Shanghai, China). Images were captured under a Zeiss LSM 880 confocal microscope.

For wheat germ agglutinin (WGA) staining, frozen sections (10-µm thickness) were recovered to room temperature and stained using WGA stain (W11261; ThermoFisher, Waltham, MA, USA). In brief, sections were washed with PBS twice (5 min), then fixed in 4% paraformaldehyde for 15 min at room temperature, and permeabilized with 0.5% Triton X-100 for 10 min. The sections were incubated with diluted WGA (5 μg /ml) for 60 min, washed with PBS twice, and mounted with anti-fade mounting medium. Images were captured under a Zeiss LSM 880 confocal microscope. Sections were washed with PBS twice (5 min) and mounted with neutral balsam mounting medium (E675007; BBI Life Science, Shanghai, China). Images were captured under a Zeiss LSM 880 confocal microscope. For acetylcholinesterase (AChE) staining, frozen sections (6-µm thickness) were recovered to room temperature, and stained using AChE Staining Kit (G2110; Solarbio, Beijing, China). Sections were washed with PBS twice (5 min) and mounted with neutral balsam mounting medium (E675007; BBI Life Science, Shanghai, China). Images were captured under a Zeiss LSM 880 confocal microscope.

### Cell culture and transfection

Cells were maintained in Dulbecco’s modified Eagle’s medium (DMEM) (Corning, Inc., Corning, NY, USA) supplemented with 10% FBS (Pan Seratech, Aidenbach, Germany) and 1% penicillin/streptomycin solution (Gibco, Waltham, MA, USA) at 37°C with 5% CO_2_. HUVECs were maintained in endothelial cell medium (ECM, ScienCell) consisting of 500 ml basal medium, 25 ml FBS, 5 ml endothelial cell growth supplement (ECGS) and 5 ml penicillin and streptomycin (P/S) solution. Human embryonic kidney 293T (HEK293T) cells and HUVECs were obtained from ATCC (Manassas, VA, USA). Murine cardiac fibroblasts from *Ythdf3*^-/-^ and wild-type mice were isolated as described in the literature^38^. Briefly, the hearts of neonatal mice were cut out with sterile scissors and quickly placed in ice-cold sterile 1 × PBS. The hearts were transferred to 1 mL of digestion buffer (0.4 mg per ml collagenase type II and 0.25% trypsin in 1 × PBS), cut into ∼1 mm^3^ pieces, and maintained at 37°C in a shaker incubator for 10 min with pipetting up and down 4-8 times. The digestion solution was collected into 2 mL horse serum (Meilunbio, Dalian, China), seeded into 6-well plates (one heart in each well), and incubated in a CO_2_ incubator for 1 h to allow the attachment of fibroblasts. The supernatant was removed, and the cells were cultured in DMEM at 37°C with 5% CO_2_ for further analysis.

For RNA interference, siRNA (see **Table S3** for sequences) was used to transfect cells using Lipofectamine RNAiMAX (13778150; ThermoFisher, Waltham, MA, USA), following the manufacturer’s instruction. All siRNAs were purchased from GenePharma (Suzhou, China). Plasmid transfection was performed using Lipofectamine 3000 (L3000015; ThermoFisher), following the manufacturer’s instructions. For lentiviral transfection, we co-transfected pCDH plasmids encompassing the target sequences (9 μg), or pLKO.1-shRNA (9 μg), psPAX2 (6 μg), and pMD2G (3 μg) plasmids, into 10-cm dishes of cultured HEK293T cells at 70% confluence. After 48 h of transfection, the cell supernatants were collected and filtered through 0.45-µm membranes (Merck, Darmstadt, Germany). The filtered supernatant was added to the target cells, and 48 h later, 2 μg/ml puromycin was added and incubated for 2 days to select the stable cell line.

### Immunofluorescence microscopy

Cells were seeded onto glass coverslips in 24-well plates (WHB-24-CS; WHB, Shanghai, China) and cultured until 40% confluence. Frozen sections (10-µm thickness) of mouse tissues and cells on coverslips were washed with PBS twice (5 min), then fixed in 4% paraformaldehyde (PFA) for 15 min at room temperature. Samples were permeabilized with 0.5% Triton X-100 for 10 min, blocked with 5% bovine serum albumin (1 h), and incubated with diluted primary antibody at 4°C overnight, followed by two washes of PBS (5 min). Alexa Fluor-conjugated secondary antibody was added, incubated at room temperature for 1 h, then washed with PBS twice (5 min). Slides were counterstained with 4′,6-diamidino-2-phenylindole (DAPI) (C1002; Beyotime), and mounted with anti-fade mounting (BL701A; Biosharp, Hefei, China). Images were then captured under a DragonFly confocal imaging system and analyzed with Imaris Viewer software (Andor, Belfast, UK). Images were then captured under High Intelligent and Sensitive SIM (HIS-SIM) and analyzed with HIS-SIM software (CSR Biotech, China).

### Immunoblotting

Cells were washed twice with chilled PBS buffer, scratched into RIPA buffer (MP015, Macgene, Beijing, China) and protease inhibitors, and spun at 15,000 × *g* for 30 min. The supernatant was collected, and protein concentration was determined using a BCA assay kit (23225; ThermoFisher). The proteins were resolved by SDS-PAGE and transferred to PVDF membrane (IPVH00010; Merck, Darmstadt, Germany). The membranes were blocked with 5% skimmed milk in 1 × TBST (T1082; Solarbio, Beijing, China) at room temperature for 1 h and then incubated with primary antibody at 4°C overnight. The membranes were washed three times (10 min each) in 1 × TBST at room temperature, then incubated with horseradish peroxidase-conjugated secondary antibodies at room temperature for 1 h. After washing 3 times (10 min each) in 1 × TBST at room temperature, membranes were exposed with SuperSignal West Pico PLUS Chemiluminescent Substrate (34577; ThermoFisher) in BioRad Imaging System (Hercules, CA, USA).

### RNA extraction and analysis

Cells and tissues were lysed in TRIzol reagent RNAiso PLUS (9109; TaKaRa) to extract total RNA following the manufacturer’s instruction. The extracted RNA was reverse transcribed into complementary DNA by using 5× Evo M-MLV RT Master Mix (AG11706; Accurate Biology, Changsha, China). The resulting cDNA was subjected to real-time PCR using Hieff qPCR SYBR Green Master Mix (11201ES08; Yeasen, Shanghai, China) on qTOWER (Analytik Jena, Germany) and QuantStudio (Invitrogen, MA, USA), following the manufacturer’s instructions, with the gene-specific primers listed in **Table 3**.

### RNA immunoprecipitation

RNA-IP was performed as previously described^39^. In brief, cells were cultured until an 80% confluence was achieved. Ultraviolet radiation crosslinking (400 mJ/cm²) was performed, followed by washing with PBS twice (5 min each). Cells were collected in 1.5-ml tubes and centrifuged at 600 × *g* for 5 min. They were then resuspended in 500 µl lysis buffer (50 mM HEPES pH 7.5, 150 mM KCl, 0.5% NP-40, 2mM EDTA, 0.5 mM DTT) containing protease and RNase inhibitors. Antibodies conjugated to protein A/G magnetic beads were added and incubated at 4°C overnight. This was followed by three washes with wash buffer. Then, 0.02 U DNase I (EN401; Vazyme, Nanjing, China) was added to 100 µl NT2 buffer (50 mM Tris-HCl pH 7.5, 150 mM NaCl, 1 mM MgCl_2_, 0.05% NP-40) and incubated at 30°C for 15 min. Subsequently, the supernatant was discarded, 5 μl proteinase K (ST532; Beyotime) was added into 100 µl NT2 buffer and incubated at 55°C for 15 min. The supernatant was collected into TRIzol, followed by RNA extraction and RT-qPCR.

### RNA-fluorescence in situ hybridization

Cells were carefully seeded onto glass coverslips placed within 24-well plates and cultured until a 40% confluence was attained. The cells were thoroughly washed with PBS twice, with each wash lasting for 5 min, followed by fixation in 4% paraformaldehyde for 15 min at room temperature. Subsequently, they were permeabilized with 0.1% Triton X-100 for 10 min and rewashed with PBS twice, each for 5 min. The RNA FISH SA-Biotin kit (GenePharma, China) was employed in accordance with the manufacturer’s instructions. Regarding the sequences, please refer to **Table S3**. Briefly, 1 × blocking solution was added to each well and incubated at 37°C for 30 min, followed by washing with 2 × SCC buffer at 37°C for 30 min. Probe buffer was added to each well and incubated at 37°C for 12-16 h. Then, 1 μL of 1 μM probe-biotin (denatured at 75°C for 10 min), 1 μL of 1 μM SA-Cy3, and 9 μL 1 × PBS were incubated at 37°C for 30 min and then mixed with 90 μL Buffer E from this kit to form the probe buffer. SA-Cy3 (streptavidin protein) combines with probe-biotin, where each SA can combine with four oligo-biotin and has six Cy3 on each SA, indicating that each RNA connection involves 24 Cy3. After 12-16 h, they were washed with 0.1% Buffer F (from the kit) at 37°C for 10 min, washed with 2 × SCC buffer at 60°C for 10 min twice, and washed with 2 × SCC buffer at 37°C for 10 min twice, then antibody incubation was carried out. The subsequent steps were as described for the immunofluorescence analysis. Images were captured via a DragonFly confocal imaging system (Andor) and analyzed using Imaris Viewer software (Andor).

### MS2 and MS2-coat protein (MCP) imaging system

Green fluorescent protein (GFP) was fused to the RNA-binding protein MS2 (MCP). A nuclear-localization signal was engineered into this GFP-MS2. Cells were seeded onto glass coverslips placed within 24-well plates and cultured until a 40% confluence. The lentivirus carrying Sox9-MS2 loops and MCP-GFP was added to the cells. After 48 h, direct observation or immunofluorescence was performed.

### mRNA stability

Cells were cultivated until they reached 80% confluence within a 6-well plate. One well was harvested as the sample at 0 h. To the remaining wells, we incorporated actinomycin D stock to achieve a final concentration of 10 µg/ml in culture media. Samples were gathered at diverse time points subsequent to the addition of actinomycin D, followed by RNA extraction and RT-qPCR.

### Protein stability

Cells were cultivated until 80% confluence in 6-well plates. Some cells were harvested as the sample at 0 h. For the remaining, cycloheximide (CHX) was added to achieve a final concentration of 50 µg/ml, and samples were collected at different time points. In some case, CHX and chloroquine (CQ, inhibitor of lysosome degradation) mixture was added; in another case, CHX and MG132 (inhibition of proteasome degradation) were added. Total cell lysate was collected and subjected for immunoblotting.

### Statistical analysis and reproducibility

Statistical analysis was conducted using GraphPad Prism 7 software (GraphPad, La Jolla, CA, USA). All data are presented as mean ± standard deviation or mean ± standard error of the mean. Unpaired Student’s *t*-tests were employed for significance analysis. *P* value < 0.05 was regarded as statistically significant. For *in vivo* animal studies, at least five samples were included, and random assignment was used to form the different experimental groups. *In vitro* analysis involved at least three repetitions. No animals or data points were excluded for any reason. The experimenters were kept blind to the group assignments in all mouse experiments.

## Author contribution

C.Y. and B.L. designed the overall experiments; C.Y., C.X., J.Z., and M.W. performed most of the experiments; C.Y. and M.Q. analyzed the data; C.Y. and B.L. did the omics data analysis; C.Y. and B.L. organized the data and wrote the manuscript.

## Acknowledgments

This study was supported by the grants from the National Natural Science Foundation of China (grant nos. 82488301, 82125012, and 32430048 to B.L.), the National Key R&D Program of China STI2030-Major Projects (grant no. 2021ZD0202400 to B.L.), the Shenzhen Municipal Commission of Science and Technology Innovation (grant nos. JCYJ20220818100016035 and JCYJ20220815150210001 to B.L.). The authors would like to thank Dr. Jessica Tamanini (Shenzhen University and ETediting) for editing the manuscript prior to submission.

## Reference

1 Hong, K. N. et al. International Consensus on Differential Diagnosis and Management of Patients With Danon Disease: JACC State-of-the-Art Review. J Am Coll Cardiol 82, 1628–1647 (2023). 10.1016/j.jacc.2023.08.014

2 D’Souza R, S. et al. Danon disease: clinical features, evaluation, and management. Circ Heart Fail 7, 843–849 (2014). 10.1161/CIRCHEARTFAILURE.114.001105

3 Endo, Y., Furuta, A. & Nishino, I. Danon disease: a phenotypic expression of LAMP-2 deficiency. Acta Neuropathol 129, 391–398 (2015). 10.1007/s00401-015-1385-4

4 Nishino, I. et al. Primary LAMP-2 deficiency causes X-linked vacuolar cardiomyopathy and myopathy (Danon disease). Nature 406, 906–910 (2000). Doi 10.1038/35022604

5 Zhai, Y. et al. Clinical features of Danon disease and insights gained from LAMP-2 deficiency models. Trends Cardiovasc Med 33, 81–89 (2023). 10.1016/j.tcm.2021.10.012

6 Perez, L. et al. LAMP-2C Inhibits MHC Class II Presentation of Cytoplasmic Antigens by Disrupting Chaperone-Mediated Autophagy. J Immunol 196, 2457–2465 (2016). 10.4049/jimmunol.1501476

7 Manso, A. M. et al. Systemic AAV9.LAMP2B injection reverses metabolic and physiologic multiorgan dysfunction in a murine model of Danon disease. Sci Transl Med 12 (2020). 10.1126/scitranslmed.aax1744

8 Wang, X. & He, C. Dynamic RNA modifications in posttranscriptional regulation. Mol Cell 56, 5–12 (2014). 10.1016/j.molcel.2014.09.001

9 Yang, Y., Hsu, P. J., Chen, Y. S. & Yang, Y. G. Dynamic transcriptomic m(6)A decoration: writers, erasers, readers and functions in RNA metabolism. Cell Res 28, 616–624 (2018). 10.1038/s41422-018-0040-8

10 Dorn, L. E. et al. The N(6)-Methyladenosine mRNA Methylase METTL3 Controls Cardiac Homeostasis and Hypertrophy. Circulation 139, 533–545 (2019). 10.1161/CIRCULATIONAHA.118.036146

11 Yoon, K. J. et al. Temporal Control of Mammalian Cortical Neurogenesis by m(6)A Methylation. Cell 171, 877–889.e817 (2017). 10.1016/j.cell.2017.09.003

12 Petrosino, J. M. et al. The m(6)A methyltransferase METTL3 regulates muscle maintenance and growth in mice. Nat Commun 13, 168 (2022). 10.1038/s41467-021-27848-7

13 Cao, X. et al. Mettl14-Mediated m(6)A Modification Facilitates Liver Regeneration by Maintaining Endoplasmic Reticulum Homeostasis. Cell Mol Gastroenterol Hepatol 12, 633–651 (2021). 10.1016/j.jcmgh.2021.04.001

14 Li, H. B. et al. m(6)A mRNA methylation controls T cell homeostasis by targeting the IL-7/STAT5/SOCS pathways. Nature 548, 338–342 (2017). 10.1038/nature23450

15 Wu, Y. et al. Mettl3-mediated m(6)A RNA methylation regulates the fate of bone marrow mesenchymal stem cells and osteoporosis. Nat Commun 9, 4772 (2018). 10.1038/s41467-018-06898-4

16 Golubeva, V. A. et al. Loss of YTHDF2 Alters the Expression of m(6)A-Modified Myzap and Causes Adverse Cardiac Remodeling. JACC Basic Transl Sci 8, 1180–1194 (2023). 10.1016/j.jacbts.2023.03.012

17 Li, M. et al. Ythdf2-mediated m(6)A mRNA clearance modulates neural development in mice. Genome Biol 19, 69 (2018). 10.1186/s13059-018-1436-y

18 Gilbert, C. J. et al. YTHDF2 governs muscle size through a targeted modulation of proteostasis. Nat Commun 15, 2176 (2024). 10.1038/s41467-024-46546-8

19 Xu, C. et al. YTHDF1 differentiates the contributing roles of mTORC1 in aging. Molecular Cell 10.1016/j.molcel.2025.05.003

20 Li, W. et al. Comprehensive analysis of RNA m6A methylation in pressure overload-induced cardiac hypertrophy. BMC Genomics 23, 576 (2022). 10.1186/s12864-022-08833-w

21 Flamand, M. N. & Meyer, K. D. m6A and YTHDF proteins contribute to the localization of select neuronal mRNAs. Nucleic Acids Res 50, 4464–4483 (2022). 10.1093/nar/gkac251

22 Wang, C. et al. FDRdb: a manually curated database of fibrotic disease-associated RNAome and high-throughput datasets. Database (Oxford) 2022 (2022). 10.1093/database/baac095

23 Chang, G. et al. YTHDF3 Induces the Translation of m(6)A-Enriched Gene Transcripts to Promote Breast Cancer Brain Metastasis. Cancer Cell 38, 857–871.e857 (2020). 10.1016/j.ccell.2020.10.004

24 Eggers, S., Ohnesorg, T. & Sinclair, A. Genetic regulation of mammalian gonad development. Nat Rev Endocrinol 10, 673–683 (2014). 10.1038/nrendo.2014.163

25 Lacraz, G. P. A. et al. Tomo-Seq Identifies SOX9 as a Key Regulator of Cardiac Fibrosis During Ischemic Injury. Circulation 136, 1396–1409 (2017). 10.1161/circulationaha.117.027832

26 Li, G., Fang, Y., Xu, N., Ding, Y. & Liu, D. Fibroblast-like synoviocytes-derived exosomal circFTO deteriorates rheumatoid arthritis by enhancing N6-methyladenosine modification of SOX9 in chondrocytes. Arthritis Res Ther 26, 56 (2024). 10.1186/s13075-024-03290-0

27 Zhou, Y., Zeng, P., Li, Y. H., Zhang, Z. & Cui, Q. SRAMP: prediction of mammalian N6-methyladenosine (m6A) sites based on sequence-derived features. Nucleic Acids Res 44, e91 (2016). 10.1093/nar/gkw104

28 Bertrand, E. et al. Localization of ASH1 mRNA particles in living yeast. Mol Cell 2, 437–445 (1998). 10.1016/s1097-2765(00)80143-4

29 Feng, Y. L., Chen, D. Q., Vaziri, N. D., Guo, Y. & Zhao, Y. Y. Small molecule inhibitors of epithelial-mesenchymal transition for the treatment of cancer and fibrosis. Med Res Rev 40, 54–78 (2020). 10.1002/med.21596

30 Jakob, S. & Lovell-Badge, R. Sex determination and the control of Sox9 expression in mammals. Febs j 278, 1002–1009 (2011). 10.1111/j.1742-4658.2011.08029.x

31 Hanley, K. P. et al. Ectopic SOX9 mediates extracellular matrix deposition characteristic of organ fibrosis. J Biol Chem 283, 14063–14071 (2008). 10.1074/jbc.M707390200

32 Aggarwal, S. et al. SOX9 switch links regeneration to fibrosis at the single-cell level in mammalian kidneys. Science 383, eadd6371 (2024). 10.1126/science.add6371

33 Trogisch, F. A. et al. Endothelial cells drive organ fibrosis in mice by inducing expression of the transcription factor SOX9. Sci Transl Med 16, eabq4581 (2024). 10.1126/scitranslmed.abq4581

34 Pask, A. J., Calatayud, N. E., Shaw, G., Wood, W. M. & Renfree, M. B. Oestrogen blocks the nuclear entry of SOX9 in the developing gonad of a marsupial mammal. BMC Biol 8, 113 (2010). 10.1186/1741-7007-8-113

35 Stewart, M. K., Mattiske, D. M. & Pask, A. J. Estrogen suppresses SOX9 and activates markers of female development in a human testis-derived cell line. BMC Mol Cell Biol 21, 66 (2020). 10.1186/s12860-020-00307-9

36 Sun, J. et al. A Glb1-2A-mCherry reporter monitors systemic aging and predicts lifespan in middle-aged mice. Nat Commun 13, 7028 (2022). 10.1038/s41467-022-34801-9

37 Iglesias-Pedraz, J. M. & Comai, L. Measurements of Hydrogen Peroxide and Oxidative DNA Damage in a Cell Model of Premature Aging. Methods Mol Biol 2144, 245–257 (2020). 10.1007/978-1-0716-0592-9_22

38 Kumar, S., Nagesh, D., Ramasubbu, V., Prabhashankar, A. B. & Sundaresan, N. R. Isolation and Culture of Primary Fibroblasts from Neonatal Murine Hearts to Study Cardiac Fibrosis. Bio Protoc 13, e4616 (2023). 10.21769/BioProtoc.4616

39 Xu, C. et al. HnRNP F/H associate with hTERC and telomerase holoenzyme to modulate telomerase function and promote cell proliferation. Cell Death Differ 27, 1998–2013 (2020). 10.1038/s41418-019-0483-6

